# Synthetic metabolic pathways for ethylene glycol assimilation outperform natural counterparts

**DOI:** 10.1101/2024.09.05.611552

**Authors:** Michelle Feigis, Radhakrishnan Mahadevan

## Abstract

Biomanufacturing can play a pivotal role in the transition away from fossil fuel dependence for the production of chemicals and fuels. There is growing interest in alternative bioproduction feedstocks to conventional sugars that do not compete for land use with food production. Ethylene glycol, a C2 compound that can be recovered from plastic waste or derived from CO_2_ with increasing efficiency, is gaining attention as a carbon source for microbial processes. Here we review the natural and synthetic metabolic pathways currently available for ethylene glycol assimilation. The pathways are compared in terms of their maximum theoretical yields for biomass and value-added products, thermodynamic favourability, minimum enzyme costs, and orthogonality to central carbon metabolism. We find that synthetic pathways outperform their natural counterparts in terms of higher thermodynamic driving forces, reduced enzyme costs, and higher theoretical yields for the majority of bioproducts analyzed as well as for biomass. However, natural assimilation pathways are equally or even more orthogonal to growth-associated reactions than synthetic pathways. Given these tradeoffs, the optimal EG assimilation pathway may depend on product and process choice.

## 1 Introduction

Rising concerns about climate change and anthropogenic environmental issues are motivating efforts to shift away from fossil fuel dependence in the energy and chemical production sectors. While biomanufacturing has the potential to significantly shift chemical production away from reliance on fossil fuels and help curb carbon emissions, most industrial applications thus far have been at the relatively small volumes needed for pharmaceutical and specialty chemical markets (Scown, 2022). The widespread adoption of bioproduction for bulk chemicals produced at very large scales is currently hindered by the high cost associated with microbial fermentation, much of which arises from the carbon source, conventionally simple sugars such as glucose derived from biomass (Burg et al., 2016; Chen, 2012). The cultivation of feedstock biomass requires extensive land use, directly competing with the food and feed industries and raising concerns about responsible land and resource management (Muscat et al., 2020; Harvey and Pilgrim, 2011). As a result, there is growing interest in developing bioprocesses based on renewable and inexpensive alternative carbon sources, ideally those that can be derived from waste streams or industrially emitted CO_2_.

Ethylene glycol (EG), a C2 compound and the simplest diol, is an industrial chemical readily available at large scales for relatively low cost. Although commonly known for its use as an antifreeze agent, the use of EG is widespread among many industries involved with energy, chemical synthesis, textiles, automotives and manufacturing technologies (Yue et al., 2012). EG is a component and degradation product of polyethylene terephthalate (PET), a ubiquitous polymer used for short-lived products such as single-use plastic bottles and packaging, fabrics, and textiles, with a production volume of approximately 70 million tons per year (Neves Ricarte et al., 2021). The recycling of PET is not currently considered profitable due to its low cost relative to the recycling process cost, raising environmental concerns regarding PET waste management (Sheldon and Norton, 2020). This has prompted interest in the upcycling of both its EG and terephthalic acid (TPA) monomer constituents into value-added products. PET degradation has been demonstrated via various chemical hydrolysis methods (Cao et al., 2022; Gao et al., 2022; Panda et al., 2021), enzymatic depolymerization (Ellis et al., 2021; Tiso et al., 2022; Erickson et al., 2022; Kosiorowska et al., 2022), and in vivo biological degradation using microbial hosts (Tiso et al., 2021; Brandenberg et al., 2022; Liu et al., 2022; Yoshida et al., 2016).

Aside from plastic waste, EG can be generated from such renewable sources as CO_2_ (Fan et al., 2023; Lum et al., 2020), syngas (Gor et al., 2023; Tremblay, 2011), and lignocellulosic biomass (Enjamuri and Darbha, 2022; Tullo, 2012). While the direct electrochemical reduction of CO_2_ to EG is possible via copper (Kuhl et al., 2012) or goldbased electrodes (Tamura et al., 2015), the technology is at an early stage of development. More commonly, CO_2_ is converted to ethylene (Prajapati et al., 2022; Leow et al., 2020; Nam et al., 2022) or ethylene oxide (Y. Li et al., 2022) which can be further oxidized to EG, an area that has seen recent improvements in terms of achievable current densities and product selectivity (Fan et al., 2023; A.-Z. Li et al., 2024; Lum et al., 2020).

While other C1 and C2 compounds can be generated from CO_2_ via electrochemical reduction, such as methanol, formate, ethanol, and acetate, the use of EG as a microbial feedstock for bioproduction has been comparatively less explored. In terms of physical properties, it is particularly favourable for such use as it is a relatively non-volatile, low viscosity liquid that is completely soluble in aqueous media (Yue et al., 2012). The conversion of EG to value-added products has been demonstrated in both *Escherichia coli* and *Pseudomonas putida* with the production of glycolic acid (Pandit et al., 2021), aromatic amino acids (Panda et al., 2023), and 2,4-dihydroxybutyric acid (Frazão et al., 2023).

Compared to methanol, another alternative (C1) feedstock garnering attention, EG assimilation involves the formation of less toxic intermediates (glycolaldehyde instead of formaldehyde), a lower O_2_ demand for its bioconversion to many products, and physicochemical properties that create safer conditions in the context of industrial fermentation (Wagner, Wen, et al., 2023). Both natural and synthetic pathways for the assimilation of EG into microbial metabolism have been discovered, conceptualized, and/or implemented to various degrees. Here, we provide an overview of these pathways and evaluate their amenability for bioproduction based on theoretical metrics. We look at the maximum biomass and product yields that can be achieved with each pathway in addition to their thermodynamic feasibility, minimum enzyme costs, and orthogonality to central metabolism.

## 2 Metabolic pathways for ethylene glycol assimilation

### 2.1 Natural ethylene glycol assimilation pathways

Three types of natural EG degradation pathways are known to exist in microbial metabolism. One is primarily found in acetogens such as *Clostridium glycolicum* and *Acetobacterium woodii* and involves the the dehydration of EG to acetaldehyde using an extremely O_2_-sensitive diol-dehydratase (Hartmanis and Stadtman, 1986; Trifunovíc et al., 2016). An acetaldehyde dehydrogenase enzyme then activates acetaldehyde to acetyl-CoA, a key central carbon metabolite that serves as the entry point of the tricarboxylic acid (TCA) cycle. In aerobic organisms, all known EG assimilation routes involve the initial oxidation of EG to glycolaldehyde (GA). In the naturally-occurring ‘glycerate pathway’, GA is subsequently oxidized to glyoxylate and assimilated into central carbon metabolism via transformation to 2-phosphoglycerate of lower glycolysis or through the glyoxylate shunt (Pandit et al., 2021; Boronat et al., 1983). In *E. coli*, the initial oxidation of EG is mediated by an NAD^+^-dependent propanediol oxidoreductase (FucO) enzyme with promiscuous activity towards EG. In *Pseudomonas putida* and *Ideonella sakaiensis*, this oxidation has been found to be facilitated by pyrroloquinoline quinone (PQQ)-dependent alcohol dehydrogenases (Mückschel et al., 2012, Hachisuka et al., 2022). The conversion of EG to value-added products via the glycerate pathway has been demonstrated in *E. coli* (Pandit et al., 2021; Panda et al., 2023; Frazão et al., 2023), *P. putida* (Franden et al., 2018; Bao et al., 2023) and *Rhodococcus jostii* (Diao et al., 2023). A major drawback of this EG assimilation route is the carbon loss associated with the condensation of two glyoxylate molecules, during which 25% of the initial carbon is released as CO_2_ (Figure 2.1).

An alternative route for the assimilation of glyoxylate (an intermediate compound in the glycerate pathway) is the β-hydroxyaspartate cycle (BHAC), which was recently fully characterized in the bacteria *Paracoccus denitrificans* (Schada von Borzyskowski et al., 2019), although parts of the module were described as early as the 1960s (Kornberg and Morris, 1963; Kornberg and Morris, 1965). In this pathway, glyoxylate is added to the amino acid glycine to form (2*R*,3*S*)-β-hydroxyaspartate (BHA), which is subsequently dehydrated to iminosuccinate and reduced to L-aspartate (Figure 2.1). An aminotransferase converts L-aspartate to the TCA cycle intermediate oxaloacetate while regenerating glycine from glyoxylate. This cycle sidesteps the carbon loss and ATP requirements associated with glyoxylate assimilation in the glycerate pathway. The BHAC was recently implemented in *P. putida* KT2440, enabling it to grow on EG more efficiently in terms of growth rate and biomass yield than using the glycerate pathway (Schada von Borzyskowski et al., 2023).

### 2.2 Synthetic ethylene glycol assimilation pathways

Pathways for the assimilation of EG that do not exist in nature have also been conceptualized, and implemented in some cases, with the goal of outperforming native pathways. To the best of our knowledge, all synthetic EG assimilation pathways begin with the oxidation of EG to GA (Figure 2.1), mediated either through native aldehyde dehydrogenases such as FucO in *E. coli* or heterologous analogs with demonstrated improved activity such as Gox0313 from *Gluconobacter oxydans* (Zhang et al., 2015). In the synthetic acetyl-CoA (SACA) pathway, GA is converted to acetyl-phosphate (AcP), incorporating inorganic phosphate via an acetyl-phosphate synthase enzyme, followed by the production of acetyl-CoA from AcP by a phosphate acetyltransferase (PTA) enzyme, which is native to the *E. coli* metabolism (Lu et al., 2019). This pathway has not been shown as of yet to support microbial growth on EG.

In the tartonyl-CoA (TaCo) pathway, suggested by Scheffen et al. (Scheffen et al., 2021), GA is converted to glycolyl-CoA through an aldehyde dehydrogenase (PduP) from *Rhodopseudomonas palustris* BisB18 (Figure 2.1). A new-to-nature enzyme, glycolyl-CoA carboxylase (GCC) can covert glycolyl-CoA to (S)-tartronyl-CoA, which is finally reduced to glycerate (a central carbon metabolite) through a tartronyl-CoA reducatse (TCR). Scheffen et al. was able to demonstrate the in vitro production of glycerate from EG with simultaneous CO_2_ fixation (Scheffen et al., 2021), however this pathway has not yet been successfully implemented in vivo.

The synthetic arabinose-5-phosphate (Ara5P)-dependent glycolaldehyde assimilation (SAGA) pathway (Yang et al., 2019) provides a cyclical alternative to the SACA pathway for the production of acetyl-CoA from glycolaldehyde (Figure 2.1). An arabinose 5-phosphate aldolase (FsaA) enzyme catalyzes the addition of the acceptor molecule glyceraldehyde 3-phosphate (GA3P) to GA yielding D-arabinose 5-phsophate (Ara5P), which is rearranged to form D-ribulose 5-phosphate (Ribu5P) via an isomerase (KdsD) followed by xylulose 5-phosphate (Xylu5P) via an epimerase (Rpe). The cleavage of Xylu5P by a phosphoketolase (Pkt) regenerates GA3P while releasing acetyl phosphate (AcP), the precursor to acetyl-CoA in the reaction catalyzed by phopsphate acetyltransferase (Pta). Wagner et al. experimentally validated this pathway in *E. coli* using glycerol as the co-substrate for growth (Wagner, Bade, et al., 2023).

In a similar strategy, the synthetic allose-6-phosphate (All6P)-dependent SAGA pathway has been proposed as an analog to the Ara5P-dependent SAGA pathway, proceeding through the formation of All6P from GA and subsequent conversion to allulose-6-phosphate (Au6P), fructose-6-phosphate (F6P), and erythrose-4-phosphate (E4P), yielding AcP (Figure 2.1) (Mao et al., 2021). Mao et al. used a combinatorial algorithm combined with parsimonious flux balance analysis (pFBA) to predict eight novel carbonconserving pathways for formaldehyde assimilation, all of which can also serve as GA (and by extension, EG) assimilation pathways as they begin with the conversion of formaldehyde to GA. Of these, the All6P-dependent SAGA pathway was demonstrated in vitro (with a carbon yield of 94%) as it met the criteria of consisting of 10 or fewer reactions, exhibiting no carbon loss, being independent of ATP and reducing equivalents, and having enzymes identified to catalyze each reaction (Mao et al., 2021).

## 3 Methods

### 3.1 Theoretical yield calculations

Flux balance analysis (FBA) was performed using the COBRApy toolbox (Ebrahim et al., 2013) on Python 3.7.9 using the GLPK solver (GNU Project). The *E. coli* genome-scale model iML1515 (BiGG database) was used for all maximum theoretical yield (MTY) calculations. Reactions and metabolites that are heterologous to the *E. coli* metabolism or missing from the models (EG assimilation and non-native bioproduction pathways) were manually added, informed by reaction specifications in the literature (BioCyc). The substrate uptake rate was set to 10 mmol/gDW*·*hr while O_2_ uptake was constrained to a maximum of 20 mmol/gDW*·*hr to reflect aerobic conditions (Andersen and von Meyenburg, 1980). The lower bound of the ATP maintenance reaction was set to zero to allow for maximal yield determinations. Other parameters were unchanged from their default states based on BiGG model specifications. EG and bioproducts were assumed to passively diffuse into/out of the cell based on their sizes and charges, which was reflected in the constructed models.

To calculate the maximum theoretical biomass yield, the objective function of the metabolic model was set to maximize biomass generation (a pseudo-reaction that represents the overall cellular growth in the metabolic network). The FBA-predicted growth rate was divided by the substrate uptake rate (converted from a molar flux value to mass), as detailed in Equation 1 in which *Y_biomass_* is the biomass yield (g of biomass per g of substrate), *µ* is the growth rate (g/gDW*·*hr), 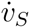 is the substrate uptake rate (mmol/gDW*·*hr), and *M_S_* is the molecular weight (g/mol) of the substrate.

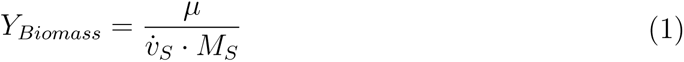

To calculate the maximum theoretical product yield, the objective function of the metabolic model was set to each bioproduct’s exchange reaction (a pseudo-reaction that represents the exchange of the product between the cytosol and extracellular matrix). The FBA-predicted product exchange fluxes were divided by the substrate uptake rate, both converted from molar flux values to mass yields, as shown in Equation 2. The product yield is denoted as *Y_Product_* (g of product per g of substrate), 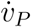 is the product exchange flux (mmol/gDW*·*hr), and *M_P_* is the molecular weight (g/mol) of the product.

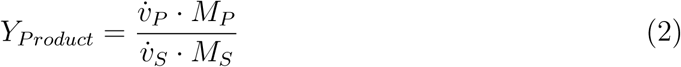

### 3.2 MDF and ECM calculations

The Max-min driving force (MDF) framework (Noor et al., 2014) was accessed through the open-source eQuilibrator API (Noor et al., 2013). The MDF provides a quantitative measure of how thermodynamically favorable a metabolic pathway is by calculating the maximum value of the minimum driving force among all reactions in the pathway. Metabolite concentrations are optimized within physiologically feasible ranges to maximize the smallest Δ*_r_G* of any reaction in the pathway. A higher value indicates a more thermodynamically favourable pathway without thermodynamic bottlenecks that is generally capable of sustaining higher fluxes (Noor et al., 2014). The MDF framework is formulated as a linear optimization problem, as shown in Equation 3, where *B* is the MDF value (kJ/mol), *S^T^* is the metabolic network represented by a stoichiometric matrix, R is the universal gas constant, T is the temperature, and x is a vector of all log values of metabolite concentrations constrained by the lower (*C_min_*) and upper (*C_max_*) bounds.

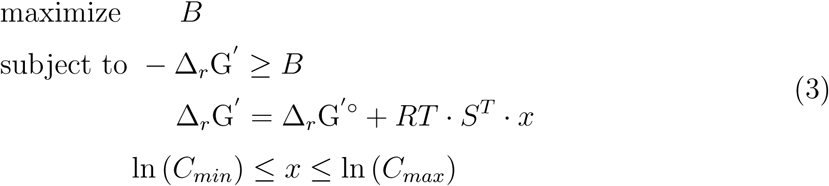

The MDF framework employs the Component Contribution method for the estimation of standard Gibbs energies of formation for each compound, from which it calculates the standard Gibbs energy change (-Δ*_r_G*′^◦^) for each reaction. A standard temperature of 25 °C was used as Gibbs energy variations with temperature are difficult to predict to match *E. coli* growth conditions (Jankowski et al., 2008). However, cellular conditions in terms of ionic strength (250 mM) (Szatmári et al., 2020) and cytosolic pH (7.5) (Slonczewski et al., 2009) were used in addition to constraining metabolite concentration to physiologically feasible values (1 *µ*M to 10 mM) (Noor et al., 2014; Bennett et al., 2009). In cases where pathways were deemed infeasible with these concentration bounds for bottleneck reactions, the upper bound of metabolite concentrations was increased (SI).

The enzyme cost minimization (ECM) framework was also accessed through the eQuilibrator API (Noor et al., 2013) and used to quantify the minimal protein cost required to support each EG assimilation pathway. Both thermodynamic and kinetic parameters are incorporated in the calculations; the same physiological conditions were used as for MDF analysis and experimental kinetic data (K*_M_* and K*_cat_*) were used wherever possible (SI).

### 3.3 Orthogonality calculations

Orthogonality is the design principle of creating independent and modular metabolic pathways that minimize interactions with the host organism’s natural cellular metabolism (Pandit et al., 2017). An orthogonality score (OS) quantifies the independence of biomass and chemical production pathways, with a higher score indicating a greater degree of separation. To calculate the orthogonality score of each pathway, *E. coli* core metabolic models (Orth et al., 2010) were used with the relevant pathway reactions added. Constraints on substrate and oxygen uptake were set as described above for genome-scale models. Elementary flux modes (EFMs), the simplest sets of reactions that can operate at steady state, were enumerated using efmtool (Terzer and Stelling, 2008) on CellNet-Analyzer for Python (CNApy; Thiele et al., 2022) for each model. EFMs were split into two distinct sets: *S_t_*, EFMs which contain a non-zero flux through the target product reaction but have zero biomass flux, and *S_X_*, EFMs with non-zero flux through the biomass reaction and zero product flux (Pandit et al., 2017). Acetate was selected as the target product as a stand-in for central biomass precursor acetyl-CoA which cannot be readily exported from the cell.

The average similarity (AS) between the reactions that are common to the EFMs in sets *S_t_* and *S_X_* is calculated using Equation 4, in which the dot product of the vectors representing these sets (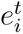 and 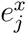, respectively) in the metabolic flux space is calculated and normalized to the size of the biomass-supporting node (Pandit et al., 2017). This value is divided by the product of m and n, the total numbers of EFMs in sets *S_t_*and *S_X_*, respectively. The OS is taken as the compliment of the AS (Equation 5).

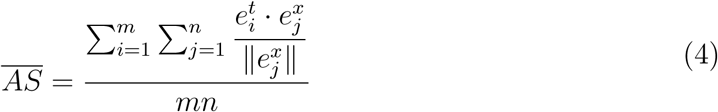

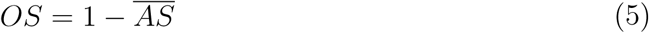

## 4 Ethylene glycol assimilation pathway evaluation

### 4.1 Theoretical yield

Metabolic pathways for EG assimilation were compared on the basis of theoretical biomass and product yields, thermodynamic favourability, enzyme burden, and orthogonality to native metabolism. Flux balance analysis (FBA) was used to calculate the theoretical biomass yield of *E. coli* with EG as the sole source of carbon and energy, in addition to a selection of bioproducts (Figure 4.1D-E) (see 3.1). In terms of theoretical yield, the synthetic pathways (SACA and SAGA) outperform the natural pathways (glycerate and BHAC) for biomass and most bioproducts, which is consistent with findings in previous analyses (Wagner, Wen, et al., 2023). Glycolate is a notable exception, for which the BHAC shows the highest yield and the glycerate pathway outperforms the SACA pathway, which is unsurprising given glycolate is an intermediate in these routes with the potential for accumulation (Pandit et al., 2021). The calculated yields for the Ara5P and All6P-dependent SAGA pathways are indistinguishable as the pathways share a common overall stoichiometry, and are consistently higher than those for the SACA pathway.

### 4.2 Thermodynamic favourability and enzyme burden

The thermodynamic favourability of each pathway up until the production of acetyl-CoA was assessed using the Max-min driving force framework (Noor et al., 2014), a tool that has been employed to assess the thermodynamic feasibility of pathways and rank competing pathways based on their overall driving force under physiological conditions (Khana et al., 2022; Shultz-Mirbach 2024). Acetyl-CoA was selected as the endpoint of each assimilation route, as it serves as the entry point towards the TCA cycle and is a crucial precursor for the production of a large number of industrially relevant compounds such as 1-butanol, fatty acids and lipids, polyketides, isoprenoids, polyhydroxyalkanoates, and and amino acids (Krivoruchko et al., 2015; Sun et al., 2020). Thermodynamically, the BHAC is the least favourable EG assimilation route (Figure 4.1A) with a negative MDF value due to thermodynamic bottlenecks in glycolate conversion to gloyxolate and its subsequent entry into the cycle via conversion to glycine and then BHA, which has a positive Δ*_r_G* value at physiological conditions (Figure 4.1; refer to SI). The SACA pathway yields the highest Max-min driving force followed by the Ara5P- and All6P-dependent SAGA pathways, suggesting a tradeoff may exist between stoichiometric yield and thermodynamic favourability (Du et al., 2018). The glycerate EG assimilation pathway has a low but positive MDF value provided that the concentration of glycolate is maintained sufficiently high in the cell (Figure 4.1A).

In order to investigate the enzyme cost to the cells of the EG assimilation pathways, the Enzyme Cost Minimization (ECM) framework (Noor et al., 2016) was used which incorporates kinetic parameters of pathway enzymes in addition to reaction thermodynamics. Unsurprisingly, the natural EG assimilation routes (glycerate pathway and BHAC) have the highest enzyme demand due to the presence of thermodynamic bottlenecks (Figure 4.1B; Figure 4.1). The significant difference in thermodynamic driving forces between the natural and synthetic pathways underlies the disparity in overall enzyme burden such that variations in kinetic parameters are negligible on the whole. The synthetic pathways are also much shorter routes to acetyl-CoA than the natural pathways, which contributes to this difference. However, as shown in Figure 4.1, kinetic constraints impose high costs of certain pathway enzymes (leading to enzyme saturation) and can limit the successful implementation of substrate assimilation pathways such as the SACA pathway, in which the key ACPS enzyme has a Michaelis-Menten constant (*K_m_*) value of 51 mM (Lu et al., 2019).

While the two SAGA pathways result in similar bioproduct yields, it is noteworthy that the All6P-dependent pathway may be more thermodynamically favourable with a lower enzyme burden than its Ara5P-dependent analog, a factor that may be considered when choosing which pathway to implement in a host organism.

### 4.3 Orthogonality

The synthesis of a desired value-added compound is often in direct competition with the natural cellular objective of maximizing growth or biomass generation, which can make achieving productivity and high chemical yields challenging. As a result, there is interest in developing bioprocesses with decoupled growth and chemical production stages, with a dynamic control scheme to switch between the two metabolic states (Ni et al., 2021; Hartline et al., 2021; Venayak et al., 2015; Raj et al., 2020). It has been suggested that orthogonal metabolic pathways, as in those that are largely independent of native metabolism with limited common metabolic nodes, may be more amenable to dynamic control compared to conventional, interconnected natural pathways such as glycolysis for glucose assimilation (Pandit et al., 2017). Indeed, our analysis using the orthogonality framework based on acetate production showed that EG assimilation pathways (with the exception of the Ara5P and All6P-dependent SAGA pathways) are more orthogonal than glucose (Figure 4.1C), with the BHAC yielding the highest orthogonality score.

## 5 Conclusions and recommendations

EG is an interesting feedstock garnering attention alongside other next-generation feedstocks such as methanol, formate, and syngas towards their microbial conversion to value-added products. The use of EG may confer advantages over such C1 compounds as methanol as its physicochemical properties are more amenable for process safety, its as-similation into microbial metabolism involves the less toxic intermediate GA instead of formaldehyde, and it can be more readily utilized as a carbon source by common industrial workhorse organisms (Wagner, Wen, et al., 2023). Recent work has focused on the implementation of EG assimilation pathways, both naturally-occurring and synthetic, in organisms such as *E. coli*, *P. putida*, and *I. sakaiensis* to produce desired target compounds. More pathways may yet be elucidated with computational tools such as bioretrosynthesis, which involves the reverse engineering of a biological pathway, starting from the desired end product and working backwards to identify necessary precursor molecules and enzymatic steps (Koch et al., 2020; Lawson et al., 2021). Furthermore, given the number of possible routes that can allow for the conversion of EG to products, the theoretical evaluation of pathways can drive experimental efforts and identify targets for further engineering.

Using an *E. coli* model, synthetic pathways (SACA and SAGA) for EG assimilation were shown to outperform natural pathways (glycerate and BHAC) in terms of higher biomass and product yields, higher thermodynamic driving forces, and lower enzyme burdens. The SACA pathway exhibits the highest thermodynamic driving force with a high orthogonality score and higher theoretical product yields than natural pathways (while lower than SAGA pathways); however its successful implementation is currently hindered by the poor kinetics of its key ACPS enzyme that converts GA to acetyl phosphate (Lu et al., 2019). Novel enzyme engineering tools, such as a synthetic orthogonal replication system (Tian et al., 2024) or machine learning-based platforms, can be used to improve enzyme performance and substrate affinities (achieve lower *K_M_* values) in such cases.

While the glycerate pathway suffers from carbon inefficiency and is not particularly advantageous in terms of thermodynamics, its relative ease of implementation and high orthogonality score may make it well-suited for the production of the pathway intermediate glycolate, especially in a two-stage system. The theoretical yield of glycolate is also higher using this pathway compared to the use of glucose or the SACA pathway. The favourability of different substrate assimilation pathways is therefore likely to be dependent on the product of interest, as has been previously suggested (Wagner, Wen, et al., 2023). Given their low orthogonality scores, the Ara5P and All6P-dependent SAGA pathways may be more suited for growth-coupled production, particularly of acetyl-CoA-derived products to which they would provide a shorter metabolic route compared to natural EG assimilation pathways.

The BHAC was identified as the EG assimilation pathway with the highest orthogonality score, suggesting it may be amenable for a two-stage bioproduction system for a product with a high theoretical yield such as glycolate. Strategies to overcome its low thermodynamic driving force could include the removal of product from the system or the selection of gaseous products to drive the pathway forward. The *E. coli* model used in this work may also not be reflective of metabolic conditions in other organisms with different metabolic networks (Nogales et al., 2020; Mo et al., 2009).

As technologies for converting CO_2_ into useful chemicals improve and the cost of these processes decreases, EG is poised to become a more appealing feedstock for bioproduction. Advances in CO_2_ electrochemical reduction may make it feasible to produce EG more efficiently and cost-effectively. It is also important to highlight that bioprocesses do not require the same level of substrate purity as chemical synthetic processes (e.g. water contamination is not a concern), which can further lower feedstock costs (Lad et al., 2022). Additionally, the ongoing development of technologies for the degradation of PET into EG further enhances the economic viability of using EG as a feedstock. As these technologies mature and scale, the reduced production costs will likely make EG a more attractive feedstock for the bioproduction of value-added chemicals.

---

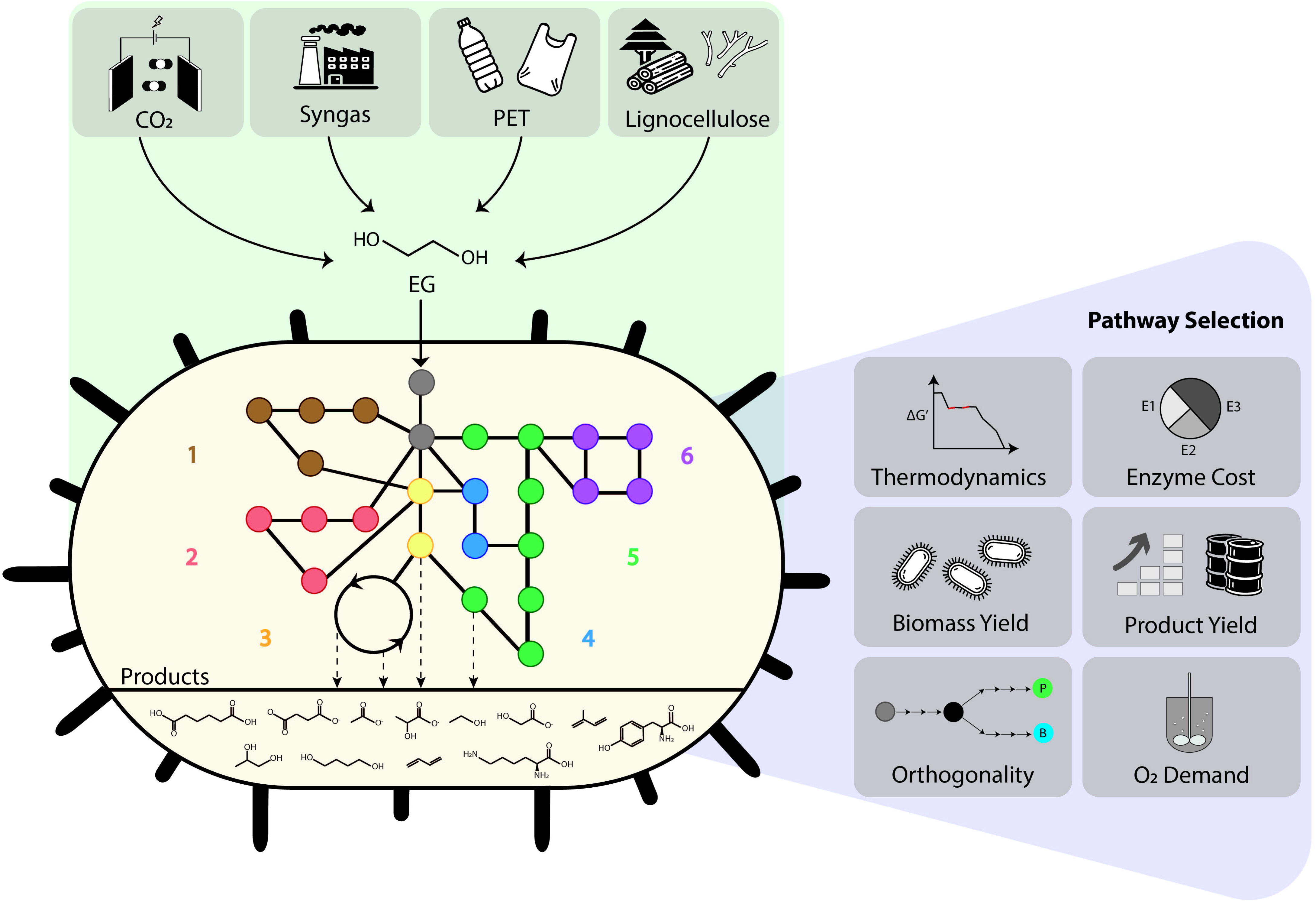

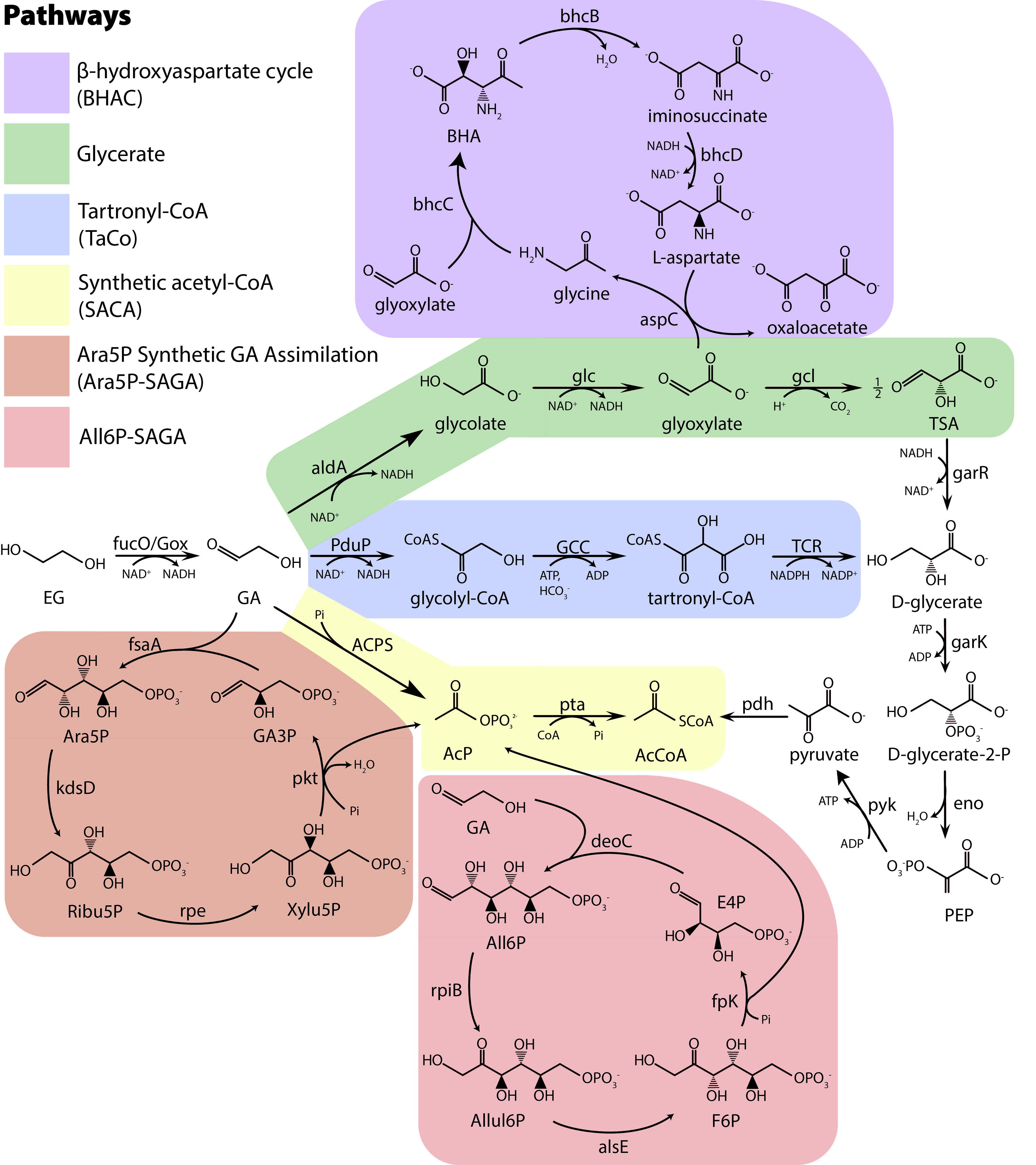

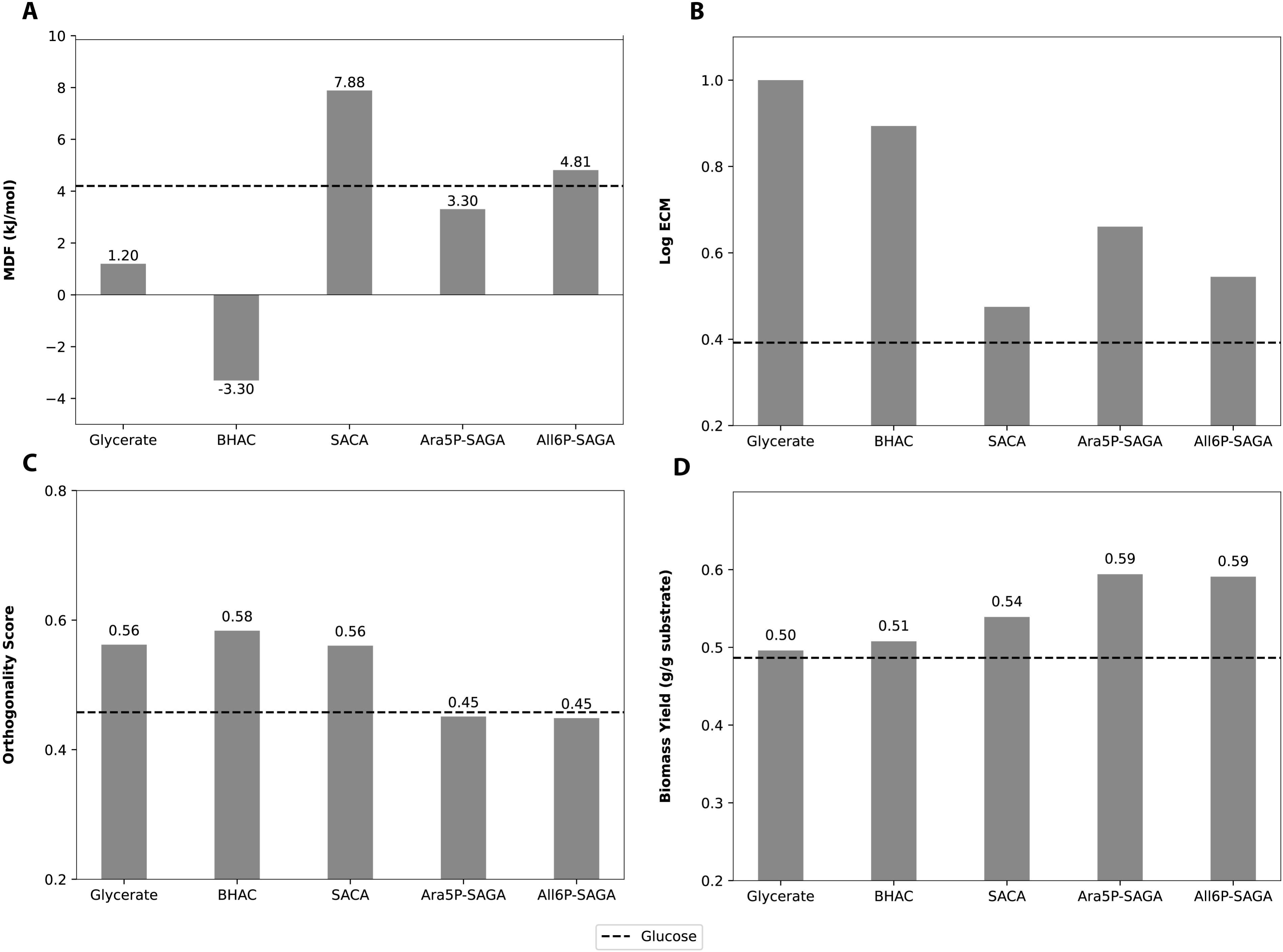

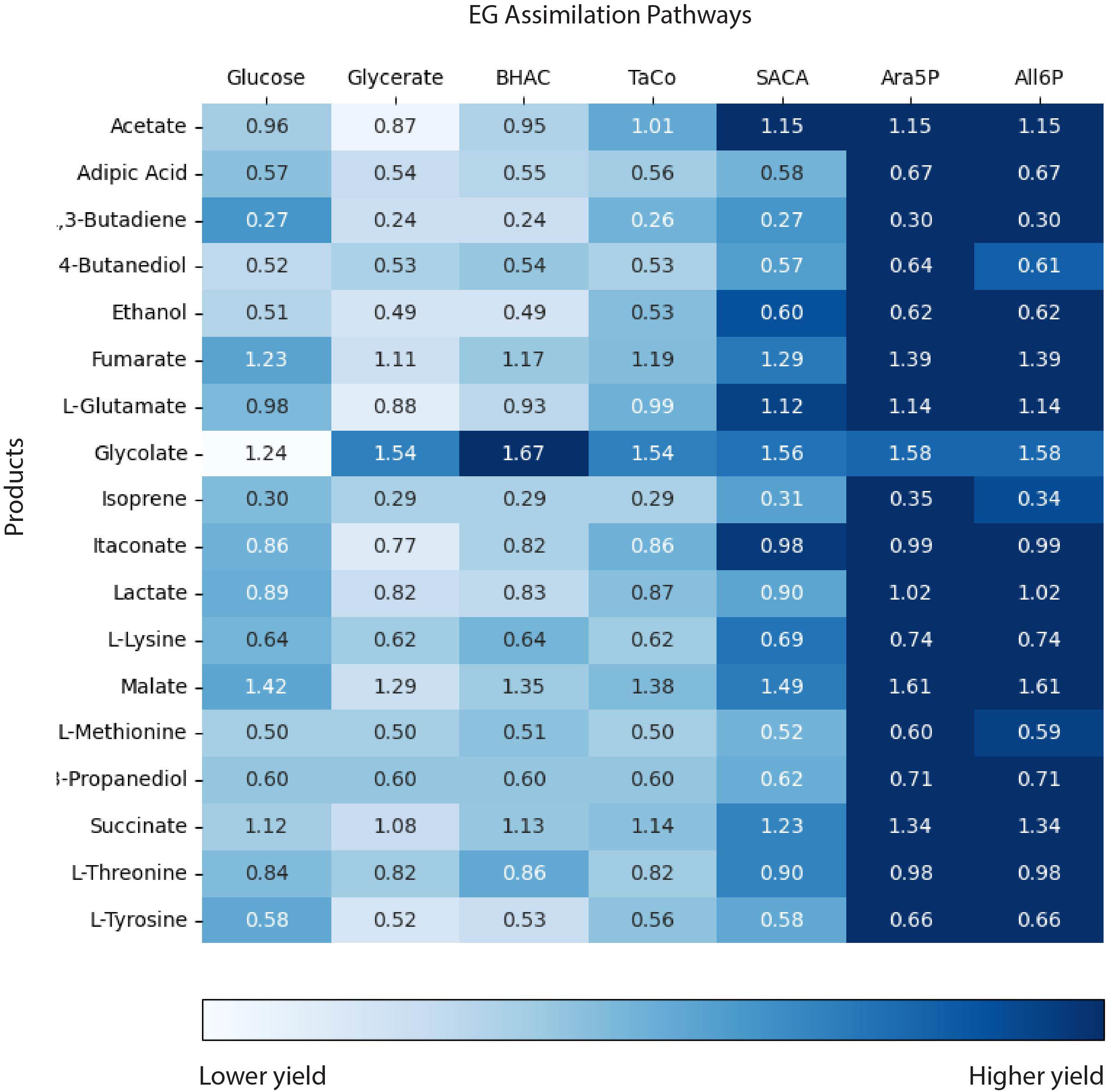

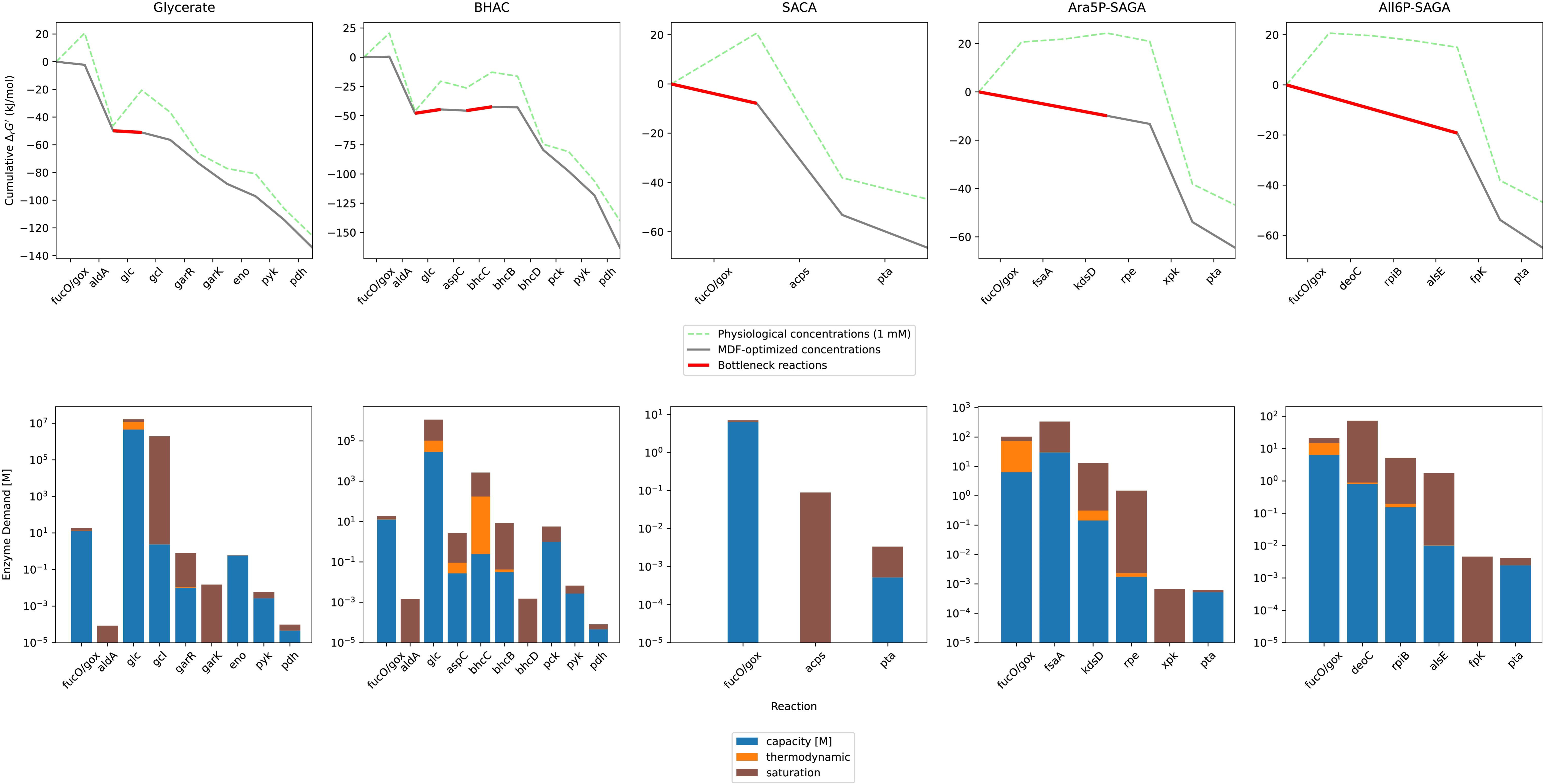

## References

1. Andersen, K. B., & von Meyenburg, K. (1980). Are growth rates of escherichia coli in batch cultures limited by respiration? Journal of Bacteriology, 144 (1), 114–123.

2. Bao, T., Qian, Y., Xin, Y., Collins, J. J., & Lu, T. (2023). Engineering microbial division of labor for plastic upcycling [Publisher: Nature Publishing Group]. Nat Commun, 14 (1), 5712.

3. Bennett, B. D., Kimball, E. H., Gao, M., Osterhout, R., Van Dien, S. J., & Rabinowitz, J. D. (2009). Absolute metabolite concentrations and implied enzyme active site occupancy in escherichia coli [Publisher: Nature Publishing Group]. Nat Chem Biol, 5 (8), 593–599.

4. Boronat, A., Caballero, E., & Aguilar, J. (1983). Experimental evolution of a metabolic pathway for ethylene glycol utilization by escherichia coli. J Bacteriol, 153 (1), 134–139.

5. Brandenberg, O. F., Schubert, O. T., & Kruglyak, L. (2022). Towards synthetic PETtrophy: Engineering pseudomonas putida for concurrent polyethylene terephthalate (PET) monomer metabolism and PET hydrolase expression. Microbial Cell Factories, 21 (1), 119.

6. Burg, J. M., Cooper, C. B., Ye, Z., Reed, B. R., Moreb, E. A., & Lynch, M. D. (2016). Large-scale bioprocess competitiveness: The potential of dynamic metabolic control in two-stage fermentations. Current Opinion in Chemical Engineering, 14, 121–136.

7. Cao, F., Wang, L., Zheng, R., Guo, L., Chen, Y., & Qian, X. (2022). Research and progress of chemical depolymerization of waste PET and high-value application of its depolymerization products [Publisher: The Royal Society of Chemistry]. RSC Adv., 12 (49), 31564–31576.

8. Chen, G.-Q. (2012). New challenges and opportunities for industrial biotechnology. Microb Cell Fact, 11 (1), 111.

9. Diao, J., Hu, Y., Tian, Y., Carr, R., & Moon, T. S. (2023). Upcycling of poly(ethylene terephthalate) to produce high-value bio-products [Publisher: Elsevier]. Cell Reports, 42 (1).

10. Du, B., Zielinski, D. C., Monk, J. M., & Palsson, B. O. (2018). Thermodynamic favorability and pathway yield as evolutionary tradeoffs in biosynthetic pathway choice. [Place: United States]. Proc Natl Acad Sci U S A, 115 (44), 11339–11344.

11. Ebrahim, A., Lerman, J. A., Palsson, B. O., & Hyduke, D. R. (2013). COBRApy: COnstraints-based reconstruction and analysis for python. BMC Systems Biology, 7 (1), 74.

12. Ellis, L. D., Rorrer, N. A., Sullivan, K. P., Otto, M., McGeehan, J. E., Román-Leshkov, Y., Wierckx, N., & Beckham, G. T. (2021). Chemical and biological catalysis for plastics recycling and upcycling [Number: 7 Publisher: Nature Publishing Group]. Nat Catal, 4 (7), 539–556.

13. Enjamuri, N., & Darbha, S. (2022). Advances in catalytic conversion of lignocellulosic biomass to ethylene glycol [Publisher: Taylor & Francis eprint: 10.1080/0161494 Catalysis Reviews, 0 (0), 1–71.

14. Erickson, E., Shakespeare, T. J., Bratti, F., Buss, B. L., Graham, R., Hawkins, M. A., König, G., Michener, W. E., Miscall, J., Ramirez, K. J., Rorrer, N. A., Zahn, M., Pickford, A. R., McGeehan, J. E., & Beckham, G. T. (2022). Comparative performance of PETase as a function of reaction conditions, substrate properties, and product accumulation. ChemSusChem, 15 (1), e202101932.

15. Fan, L., Zhao, Y., Chen, L., Chen, J., Chen, J., Yang, H., Xiao, Y., Zhang, T., Chen, J., & Wang, L. (2023). Selective production of ethylene glycol at high rate via cascade catalysis [Number: 7 Publisher: Nature Publishing Group]. Nat Catal, 6 (7), 585–595.

16. Franden, M. A., Jayakody, L. N., Li, W.-J., Wagner, N. J., Cleveland, N. S., Michener, W. E., Hauer, B., Blank, L. M., Wierckx, N., Klebensberger, J., & Beckham, G. T. (2018). Engineering pseudomonas putida KT2440 for efficient ethylene glycol utilization. Metabolic Engineering, 48, 197–207.

17. Frazão, C. J. R., Wagner, N., Rabe, K., & Walther, T. (2023). Construction of a synthetic metabolic pathway for biosynthesis of 2,4-dihydroxybutyric acid from ethylene glycol [Number: 1 Publisher: Nature Publishing Group]. Nat Commun, 14 (1), 1931.

18. Gao, Z., Ma, B., Chen, S., Tian, J., & Zhao, C. (2022). Converting waste PET plastics into automobile fuels and antifreeze components [Number: 1 Publisher: Nature Publishing Group]. Nat Commun, 13 (1), 3343.

19. Gor, N. K., Chinthala, P. K., Das, A., & Vaidya, P. D. (2023). An overview of monoethylene glycol synthesis via CO coupling reaction: Catalysts, kinetics, and reaction pathways [ eprint: https://onlinelibrary.wiley.com/doi/pdf/10.1002/cjce.24736]. The Canadian Journal of Chemical Engineering, 101 (7), 4054–4075.

20. Hachisuka, S.-i., Chong, J. F., Fujiwara, T., Takayama, A., Kawakami, Y., & Yoshida, S. (2022). Ethylene glycol metabolism in the poly(ethylene terephthalate)-degrading bacterium ideonella sakaiensis. Appl Microbiol Biotechnol, 106 (23), 7867–7878.

21. Hartline, C. J., Schmitz, A. C., Han, Y., & Zhang, F. (2021). Dynamic control in metabolic engineering: Theories, tools, and applications. Metab Eng, 63, 126–140.

22. Hartmanis, M. G. N., & Stadtman, T. C. (1986). Diol metabolism and diol dehydratase in clostridium glycolicum. Archives of Biochemistry and Biophysics, 245 (1), 144– 152.

23. Harvey, M., & Pilgrim, S. (2011). The new competition for land: Food, energy, and climate change. Food Policy, 36, S40–S51.

24. Jankowski, M. D., Henry, C. S., Broadbelt, L. J., & Hatzimanikatis, V. (2008). Group contribution method for thermodynamic analysis of complex metabolic networks. Biophysical Journal, 95 (3), 1487–1499.

25. Khana, D. B., Callaghan, M. M., & Amador-Noguez, D. (2022). Novel computational and experimental approaches for investigating the thermodynamics of metabolic networks. Current Opinion in Microbiology, 66, 21–31.

26. Koch, M., Duigou, T., & Faulon, J.-L. (2020). Reinforcement learning for bioretrosynthesis [Publisher: American Chemical Society]. ACS Synth. Biol., 9 (1), 157–168.

27. Kornberg, H. L., & Morris, J. G. (1963). Βhydroxyaspartate pathway: A new route for biosyntheses from glyoxylate [Number: 4866 Publisher: Nature Publishing Group]. Nature, 197 (4866), 456–457.

28. Kornberg, H. L., & Morris, J. G. (1965). The utilization of glycollate by micrococcus denitrificans: The β-hydroxyaspartate pathway. Biochem J, 95 (3), 577–586.

29. Kosiorowska, K. E., Moreno, A. D., Iglesias, R., Leluk, K., & Mirończuk, A. M. (2022). Production of PETase by engineered yarrowia lipolytica for efficient poly(ethylene terephthalate) biodegradation. Science of The Total Environment, 846, 157358.

30. Krivoruchko, A., Zhang, Y., Siewers, V., Chen, Y., & Nielsen, J. (2015). Microbial acetyl-CoA metabolism and metabolic engineering. Metabolic Engineering, 28, 28–42.

31. Kuhl, K. P., Cave, E. R., Abram, D. N., & Jaramillo, T. F. (2012). New insights into the electrochemical reduction of carbon dioxide on metallic copper surfaces [Publisher: The Royal Society of Chemistry]. Energy Environ. Sci., 5 (5), 7050–7059.

32. Lad, B. C., Coleman, S. M., & Alper, H. S. (2022). Microbial valorization of underutilized and nonconventional waste streams. Journal of Industrial Microbiology and Biotechnology, 49 (2), kuab056.

33. Lawson, C. E., Martí, J. M., Radivojevic, T., Jonnalagadda, S. V. R., Gentz, R., Hillson, N. J., Peisert, S., Kim, J., Simmons, B. A., Petzold, C. J., Singer, S. W., Mukhopadhyay, A., Tanjore, D., Dunn, J. G., & Garcia Martin, H. (2021). Machine learning for metabolic engineering: A review. Metabolic Engineering, 63, 34– 60.

34. Leow, W. R., Lum, Y., Ozden, A., Wang, Y., Nam, D.-H., Chen, B., Wicks, J., Zhuang, T.-T., Li, F., Sinton, D., & Sargent, E. H. (2020). Chloride-mediated selective electrosynthesis of ethylene and propylene oxides at high current density [Publisher: American Association for the Advancement of Science]. Science, 368 (6496), 1228– 1233.

35. Li, A.-Z., Yuan, B.-J., Xu, M., Wang, Y., Zhang, C., Wang, X., Wang, X., Li, J., Zheng, L., Li, B.-J., & Duan, H. (2024). One-step electrochemical ethylene-to-ethylene glycol conversion over a multitasking molecular catalyst [ eprint: 10.1021/jacs.3c14381]. Journal of the American Chemical Society, 146 (8), 5622–5633.

36. Li, Y., Ozden, A., Leow, W. R., Ou, P., Huang, J. E., Wang, Y., Bertens, K., Xu, Y., Liu, Y., Roy, C., Jiang, H., Sinton, D., Li, C., & Sargent, E. H. (2022). Redoxmediated electrosynthesis of ethylene oxide from CO2 and water. Nature Catalysis, 5 (3), 185–192.

37. Liu, P., Zheng, Y., Yuan, Y., Zhang, T., Li, Q., Liang, Q., Su, T., & Qi, Q. (2022). Valorization of polyethylene terephthalate to muconic acid by engineering pseudomonas putida [Number: 19 Publisher: Multidisciplinary Digital Publishing Institute]. International Journal of Molecular Sciences, 23 (19), 10997.

38. Lu, X., Liu, Y., Yang, Y., Wang, S., Wang, Q., Wang, X., Yan, Z., Cheng, J., Liu, C., Yang, X., Luo, H., Yang, S., Gou, J., Ye, L., Lu, L., Zhang, Z., Guo, Y., Nie, Y., Lin, J.,… Jiang, H. (2019). Constructing a synthetic pathway for acetyl-coenzyme a from one-carbon through enzyme design [Number: 1 Publisher: Nature Publishing Group]. Nature Communications, 10 (1), 1378.

39. Lum, Y., Huang, J. E., Wang, Z., Luo, M., Nam, D.-H., Leow, W. R., Chen, B., Wicks, J., Li, Y. C., Wang, Y., Dinh, C.-T., Li, J., Zhuang, T.-T., Li, F., Sham, T.-K., Sinton, D., & Sargent, E. H. (2020). Tuning OH binding energy enables selective electrochemical oxidation of ethylene to ethylene glycol [Number: 1 Publisher: Nature Publishing Group]. Nat Catal, 3 (1), 14–22.

40. Mao, Y., Yuan, Q., Yang, X., Liu, P., Cheng, Y., Luo, J., Liu, H., Yao, Y., Sun, H., Cai, T., & Ma, H. (2021). Non-natural aldol reactions enable the design and construction of novel one-carbon assimilation pathways in vitro [Publisher: Frontiers]. Front. Microbiol., 12.

41. Mo, M. L., Palsson, B. O., & Herrgård, M. J. (2009). Connecting extracellular metabolomic measurements to intracellular flux states in yeast. BMC Syst Biol, 3, 37.

42. Mückschel, B., Simon, O., Klebensberger, J., Graf, N., Rosche, B., Altenbuchner, J., Pfannstiel, J., Huber, A., & Hauer, B. (2012). Ethylene glycol metabolism by pseudomonas putida [Publisher: American Society for Microbiology]. Applied and Environmental Microbiology, 78 (24), 8531–8539.

43. Muscat, A., de Olde, E. M., de Boer, I. J. M., & Ripoll-Bosch, R. (2020). The battle for biomass: A systematic review of food-feed-fuel competition. Global Food Security, 25, 100330.

44. Nam, D.-H., Shekhah, O., Ozden, A., McCallum, C., Li, F., Wang, X., Lum, Y., Lee, T., Li, J., Wicks, J., Johnston, A., Sinton, D., Eddaoudi, M., & Sargent, E. H. (2022). High-rate and selective CO2 electrolysis to ethylene via metal–organic-framework-augmented CO2 availability [ eprint: https://onlinelibrary.wiley.com/doi/pdf/10.1002/adma.202 Advanced Materials, 34 (51), 2207088.

45. Neves Ricarte, G., Lopes Dias, M., Sirelli, L., Antunes Pereira Langone, M., Machado de Castro, A., Zarur Coelho, M. A., & Dias Ribeiro, B. (2021). Chemo-enzymatic depolymerization of industrial and assorted post-consumer poly(ethylene terephthalate) (PET) wastes using a eutectic-based catalyst [ eprint: https://onlinelibrary.wiley.com/doi/p Journal of Chemical Technology & Biotechnology, 96 (11), 3237–3244.

46. Ni, C., Dinh, C. V., & Prather, K. L. (2021). Dynamic control of metabolism [ eprint: 10.1146/annurev-chembioeng-091720-125738]. Annual Review of Chemical and Biomolecular Engineering, 12 (1), 519–541.

47. Nogales, J., Mueller, J., Gudmundsson, S., Canalejo, F. J., Duque, E., Monk, J., Feist, A. M., Ramos, J. L., Niu, W., & Palsson, B. O. (2020). High-quality genomescale metabolic modelling of pseudomonas putida highlights its broad metabolic capabilities. Environ Microbiol, 22 (1), 255–269.

48. Noor, E., Bar-Even, A., Flamholz, A., Reznik, E., Liebermeister, W., & Milo, R. (2014). Pathway thermodynamics highlights kinetic obstacles in central metabolism [Publisher: Public Library of Science]. PLOS Computational Biology, 10 (2), e1003483.

49. Noor, E., Flamholz, A., Bar-Even, A., Davidi, D., Milo, R., & Liebermeister, W. (2016). The protein cost of metabolic fluxes: Prediction from enzymatic rate laws and cost minimization [Publisher: Public Library of Science]. PLOS Computational Biology, 12 (11), e1005167.

50. Noor, E., Haraldsdóttir, H. S., Milo, R., & Fleming, R. M. T. (2013). Consistent estimation of gibbs energy using component contributions [Publisher: Public Library of Science]. PLOS Computational Biology, 9 (7), e1003098.

51. Orth, J. D., Fleming, R. M. T., & Palsson, B. Ø. (2010). Reconstruction and use of microbial metabolic networks: The core escherichia coli metabolic model as an educational guide [Company: asm Pub2Web Distributor: asm Pub2Web Institution: asm Pub2Web Label: asm Pub2Web Publisher: American Society of Microbiology]. EcoSal Plus, 4 (1).

52. Panda, S., Fung, V. Y. K., Zhou, J. F. J., Liang, H., & Zhou, K. (2021). Improving ethylene glycol utilization in escherichia coli fermentation. Biochemical Engineering Journal, 168, 107957.

53. Panda, S., Zhou, J. F. J., Feigis, M., Harrison, E., Ma, X., Fung Kin Yuen, V., Mahadevan, R., & Zhou, K. (2023). Engineering escherichia coli to produce aromatic chemicals from ethylene glycol. Metabolic Engineering, 79, 38–48.

54. Pandit, A. V., Harrison, E., & Mahadevan, R. (2021). Engineering escherichia coli for the utilization of ethylene glycol. Microbial Cell Factories, 20 (1), 22.

55. Pandit, A. V., Srinivasan, S., & Mahadevan, R. (2017). Redesigning metabolism based on orthogonality principles [Number: 1 Publisher: Nature Publishing Group]. Nature Communications, 8 (1), 15188.

56. Prajapati, A., Kani, N. C., Gauthier, J. A., Sartape, R., Xie, J., Bessa, I., Galante, M. T., Leung, S. L., Andrade, M. H. S., Somich, R. T., Rebouças, M. V., Hutras, G. T., Diniz, N., & Singh, M. R. (2022). CO2-free high-purity ethylene from electroreduction of CO2 with 4% solar-to-ethylene and 10% solar-to-carbon efficiencies [Publisher: Elsevier]. CR-PHYS-SC, 3 (9).

57. Raj, K., Venayak, N., & Mahadevan, R. (2020). Novel two-stage processes for optimal chemical production in microbes. Metabolic Engineering, 62, 186–197.

58. Schada von Borzyskowski, L., Schulz-Mirbach, H., Troncoso Castellanos, M., Severi, F., Gómez-Coronado, P. A., Paczia, N., Glatter, T., Bar-Even, A., Lindner, S. N., & Erb, T. J. (2023). Implementation of the β-hydroxyaspartate cycle increases growth performance of pseudomonas putida on the PET monomer ethylene glycol. Metabolic Engineering, 76, 97–109.

59. Schada von Borzyskowski, L., Severi, F., Krüger, K., Hermann, L., Gilardet, A., Sippel, F., Pommerenke, B., Claus, P., Cortina, N. S., Glatter, T., Zauner, S., Zarzycki, J., Fuchs, B. M., Bremer, E., Maier, U. G., Amann, R. I., & Erb, T. J. (2019). Marine proteobacteria metabolize glycolate via the β-hydroxyaspartate cycle [Number: 7783 Publisher: Nature Publishing Group]. Nature, 575 (7783), 500–504.

60. Scheffen, M., Marchal, D. G., Beneyton, T., Schuller, S. K., Klose, M., Diehl, C., Lehmann, J., Pfister, P., Carrillo, M., He, H., Aslan, S., Cortina, N. S., Claus, P., Bollschweiler, D., Baret, J.-C., Schuller, J. M., Zarzycki, J., Bar-Even, A., & Erb, T. J. (2021). A new-to-nature carboxylation module to improve natural and synthetic CO 2 fixation [Publisher: Nature Publishing Group]. Nature Catalysis, 1–11.

61. Scown, C. D. (2022). Prospects for carbon-negative biomanufacturing [Publisher: Elsevier]. Trends in Biotechnology, 40 (12), 1415–1424.

62. Sheldon, R. A., & Norton, M. (2020). Green chemistry and the plastic pollution challenge: Towards a circular economy [Publisher: The Royal Society of Chemistry]. Green Chem., 22 (19), 6310–6322.

63. Slonczewski, J. L., Fujisawa, M., Dopson, M., & Krulwich, T. A. (2009). Cytoplasmic pH measurement and homeostasis in bacteria and archaea [ISSN: 0065-2911]. In R. K. Poole (Ed.). Academic Press.

64. Sun, S., Ding, Y., Liu, M., Xian, M., & Zhao, G. (2020). Comparison of glucose, acetate and ethanol as carbon resource for production of poly(3-hydroxybutyrate) and other acetyl-CoA derivatives. Frontiers in Bioengineering and Biotechnology, 8.

65. Szatmári, D., Sárkány, P., Kocsis, B., Nagy, T., Miseta, A., Barkó, S., Longauer, B., Robinson, R. C., & Nyitrai, M. (2020). Intracellular ion concentrations and cationdependent remodelling of bacterial MreB assemblies [Publisher: Nature Publishing Group]. Sci Rep, 10 (1), 12002.

66. Tamura, J., Ono, A., Sugano, Y., Huang, C., Nishizawa, H., & Mikoshiba, S. (2015). Electrochemical reduction of CO2 to ethylene glycol on imidazolium ion-terminated self-assembly monolayer-modified au electrodes in an aqueous solution [Publisher: The Royal Society of Chemistry]. Phys. Chem. Chem. Phys., 17 (39), 26072–26078.

67. Terzer, M., & Stelling, J. (2008). Large-scale computation of elementary flux modes with bit pattern trees. Bioinformatics, 24 (19), 2229–2235.

68. Thiele, S., von Kamp, A., Bekiaris, P. S., Schneider, P., & Klamt, S. (2022). CNApy: A CellNetAnalyzer GUI in python for analyzing and designing metabolic networks. Bioinformatics, 38 (5), 1467–1469.

69. Tian, R., Rehm, F. B. H., Czernecki, D., Gu, Y., Zürcher, J. F., Liu, K. C., & Chin, J. W. (2024). Establishing a synthetic orthogonal replication system enables accelerated evolution in e. coli [Publisher: American Association for the Advancement of Science]. Science, 383 (6681), 421–426.

70. Tiso, T., Narancic, T., Wei, R., Pollet, E., Beagan, N., Schröder, K., Honak, A., Jiang, M., Kenny, S. T., Wierckx, N., Perrin, R., Avérous, L., Zimmermann, W., O’Connor, K., & Blank, L. M. (2021). Towards bio-upcycling of polyethylene terephthalate. Metabolic Engineering, 66, 167–178.

71. Tiso, T., Winter, B., Wei, R., Hee, J., de Witt, J., Wierckx, N., Quicker, P., Bornscheuer, U. T., Bardow, A., Nogales, J., & Blank, L. M. (2022). The metabolic potential of plastics as biotechnological carbon sources – review and targets for the future. Metabolic Engineering, 71, 77–98.

72. Tremblay, J.-F. (2011). Chinese firm to make ethylene glycol. Chemical & Engineering News, 89 (9).

73. Trifunovíc, D., Schuchmann, K., & Müller, V. (2016). Ethylene glycol metabolism in the acetogen acetobacterium woodii [Publisher: American Society for Microbiology]. Journal of Bacteriology, 198 (7), 1058–1065.

74. Tullo, A. (2012). Coke plays spin the bottle. Chemical & Engineering News, 90 (4).

75. Venayak, N., Anesiadis, N., Cluett, W. R., & Mahadevan, R. (2015). Engineering metabolism through dynamic control. Curr Opin Biotechnol, 34, 142–152.

76. Wagner, N., Bade, F., Straube, E., Rabe, K., Frazão, C. J. R., & Walther, T. (2023). In vivo implementation of a synthetic metabolic pathway for the carbon-conserving conversion of glycolaldehyde to acetyl-CoA. Front Bioeng Biotechnol, 11, 1125544.

77. Wagner, N., Wen, L., Frazão, C. J. R., & Walther, T. (2023). Next-generation feedstocks methanol and ethylene glycol and their potential in industrial biotechnology. Biotechnol Adv, 69, 108276.

78. Yang, X., Yuan, Q., Luo, H., Li, F., Mao, Y., Zhao, X., Du, J., Li, P., Ju, X., Zheng, Y., Chen, Y., Liu, Y., Jiang, H., Yao, Y., Ma, H., & Ma, Y. (2019). Systematic design and in vitro validation of novel one-carbon assimilation pathways. Metabolic Engineering, 56, 142–153.

79. Yoshida, S., Hiraga, K., Takehana, T., Taniguchi, I., Yamaji, H., Maeda, Y., Toyohara, K., Miyamoto, K., Kimura, Y., & Oda, K. (2016). A bacterium that degrades and assimilates poly(ethylene terephthalate) [Publisher: American Association for the Advancement of Science]. Science, 351 (6278), 1196–1199.

80. Yue, H., Zhao, Y., Ma, X., & Gong, J. (2012). Ethylene glycol: Properties, synthesis, and applications [Publisher: The Royal Society of Chemistry]. Chem. Soc. Rev., 41 (11), 4218–4244.

81. Zhang, X., Zhang, B., Lin, J., & Wei, D. (2015). Oxidation of ethylene glycol to glycolaldehyde using a highly selective alcohol dehydrogenase from gluconobacter oxydans. Journal of Molecular Catalysis B: Enzymatic, 112, 69–75.

